# Genetic fingerprinting of salmon louse (*Lepeophtheirus salmonis*) populations in the North-East Atlantic using a random forest classification approach

**DOI:** 10.1101/179218

**Authors:** A. Jacobs, M. De Noia, K. Praebel, Ø Kanstad-Hanssen, M. Paterno, D. Jackson, P. McGinnity, A. Sturm, KR Elmer, MS Llewellyn

## Abstract

Caligid sea lice represent a significant threat to salmonid aquaculture worldwide. *Lepeophtheirus salmonis* is the predominant species that occurs in the Northern Hemisphere. Dispersal of sea lice between marine aquaculture sites and geographic regions is thought to occur rapidly via planktonic transport of larvae. Population genetic analyses have consistently shown minimal population genetic structure in North Atlantic *L. salmonis*, frustrating efforts to track louse populations, improve targeted control measures and understand local adaption to environmental conditions. The aim of this study was to test the power of reduced representation library sequencing (IIb-RAD sequencing) coupled with random forest machine learning algorithms to define markers for fine-scale discrimination of louse populations. We identified 1286 robustly supported SNPs among four *L. salmonis* populations from Ireland (N=2, 27 individuals), Scotland (N=1, 11 individuals) and North Norway (N=1, 12 individuals). Weak global structure (F_SC_ = 0.018, p<0.0001) and only one significant pairwise F_ST_ comparison was observed (Scotland vs Kenmare Bay, (F_ST_ = 0.018, p<0.0001)) using all 1286 SNPs. The application of a random forest machine-learning algorithm identified 98 discriminatory SNPs that dramatically improved population assignment (DAPC assignment probability = 1), increased global F_sc_ = 0.098, (p<0.0001) and resulted in pairwise comparisons that all showed highly significant Fst-values (range = 0.081 – 0.096, p<0.0001). Out of 19 SNPs found to be under directional selection between populations, 12 corresponded to the discriminatory SNPs identified using random forest. Taken together our data suggest that *L. salmonis* SNP diversity exists with which it is possible to discriminate differences between nearby populations given suitable marker selection approaches, and that such differences might have an adaptive basis. We discuss these data in light of sea lice adaption to anthropogenic and environmental pressures as well as novel approaches to track and predict sea louse dispersal.

## Introduction

Caligid sea lice are copepod ectoparasites of marine fish. In the northern hemisphere, the salmon louse (*Lepeophtheirus salmonis*) is the species most commonly infecting farmed and wild salmonids (Boxaspen 2006), at considerable cost to animal health, biodiversity security, and economic growth. Conservative estimates of costs and losses attributed to sea louse infections, (estimated at €350M million in 2014 in Norway alone (Carmona-Antoñanzas *et al*. 2017) suggest these are the single greatest pathogen burden on the global salmonid aquaculture industry. The life cycle of the sea louse involves high levels of replication, dispersal and obligate host-association (Boxaspen 2006); this means that local environmental conditions, sea currents, and population densities are important ecological and demographic conditions to facilitate or impede infestation (Boxaspen 2006; Jackson *et al*. 2012; Salama *et al*. 2013). Eggs carried by females hatch to free-swimming non-feeding nauplii, planktonic larvae that are passively dispersed. These nauplii undergo two moults before developing into a free swimming copipodid. Development time is temperature dependent and at 10 ° C the infectious copepodid stage, which needs to settle successfully on a host for survival, develops two to three days post hatching. During the host-associated phase of the lifecycle, which progresses through further larval and preadult stages before reaching the reproducing adult stage, salmon lice feed on mucus, skin and blood of their host fish (Boxaspen 2006). Depending on severity, infections can cause skin lesions, anaemia, osmoregulatory dysfunction, stress, suppression of growth and immune function, secondary infections and, if left untreated, mortality (Boxaspen 2006; Jackson *et al*. 2017). Salmon louse control has traditionally relied on a limited number of drug treatments (Burridge *et al*. 2010; Jackson *et al*. 2017), but large-scale reliance on just a few products is associated with a significant risk of developing drug resistance (Denholm *et al*. 2002; Jackson *et al*. 2017)

Understanding and predicting salmon louse dispersal is a crucial element for predicting infestation, connectivity and the spread of salmon lice and associated drug resistance alleles. There have been several attempts to characterize population genetic structure in *L. salmonis* in the North Atlantic using conventional microsatellite and sequence markers (Glover *et al*. 2011; Nolan & Powell 2009; Tjensvoll *et al*. 2006; Todd *et al*. 1997; Todd *et al*. 2004). High gene flow between sites is consistently reported. In the largest such study (13 microsatellite loci, 2500 samples), significant but weak (0.0022) F_ST_ was detected across the Atlantic, with no evidence for population genetic structuring within geographic regions. More recently a genome-wide SNP array was developed and deployed using 6000 variable markers, and showed similar results in terms of population structure, alongside extensive evidence of selective sweeps and linkage disequilibrium attributable, at least in part, by the use of chemotherapeutics in aquaculture (Besnier *et al*. 2014). Thus, although significant progress has been made in determining population genetic signatures of selection in *L. salmonis*, the goal of distinguishing louse populations occurring in different regions - a valuable component of detecting dispersal of lice between farms – remains difficult.

Determining genetic structure in pelagic marine species has always been challenging. High rates of adult and larval dispersal impedes the accumulation of neutral variation among populations and regions. Nonetheless, several studies have achieved genetic stock delineation by focusing on non-neutral or putative adaptive markers in conjunction with high numbers of SNP markers (e.g. (Montes *et al*. 2013; Nielsen *et al*. 2012)). In extreme cases like *Anguilla rostrata*, where the organism’s reproductive ecology predicts and the genetic data support panmixia among different populations, the challenge of determining genetic differences between different populations is even greater (Côté *et al*. 2013; Jessop *et al*. 2008). Screening thousands of variable SNP markers against population genetic summary statistics may be able to detect outliers, however the identification of which markers might best assign individuals to their appropriate populations, groups, or ecomorphs necessitates further computational approaches. To this end, population genetics can usefully borrow from machine learning algorithms developed in the context of genome-wide association studies (Goldstein *et al*. 2011). Such approaches have been successfully used in *A. rostrata*, to identify SNPs that predict rearing habitat as the result of intra-generational selection (Laporte *et al*. 2016), for example. More recently similar approaches have been employed to successfully discriminate *Salmo salar* populations (Sylvester *et al*. 2017).

In this study, we identify population structure and loci under selection in *L. salmonis* using high throughput SNP genotyping and advanced analytical methods. To achieve this we collected *L. salmonis* from four different sites in the North-Eastern Atlantic (UK, Norway and two sites in Ireland) and generated genomic SNP data using a IIB restriction-enzyme associated library preparation approach (Wang *et al*. 2012). We then tested the power of Random Forest machine learning to reveal population structure and find the method reveals previously un-recognized population differences and fine-scale population differentiation.

## Methods

### Sample collection and DNA isolation

Adult and pre-adult *Lepeophtheirus salmonis* were collected in four sites around the North-East Atlantic from 18-24 month old Atlantic Salmon from commercial pens in 2015. Sites included Finnkirka (NF), Lebesby, Norway; Loch Duart (LD), Scotland, UK; Kenmare Bay (SWI), Kenmare, Ireland and Kilkieran Bay (KB), Galway, Ireland (Fig. 1A). Male and immature female individuals only were selected for sequencing to avoid gamete contamination of DNA extracts. High molecular weight DNA was obtained using a modified salt extraction protocol (See supplementary data), quantified using a NanoDrop^®^ ND-1000 spectrophotometer and visualised on a 1.5% agarose gel to assess quality. Fifteen high quality extracts were chosen per site.

**Figure 1:**
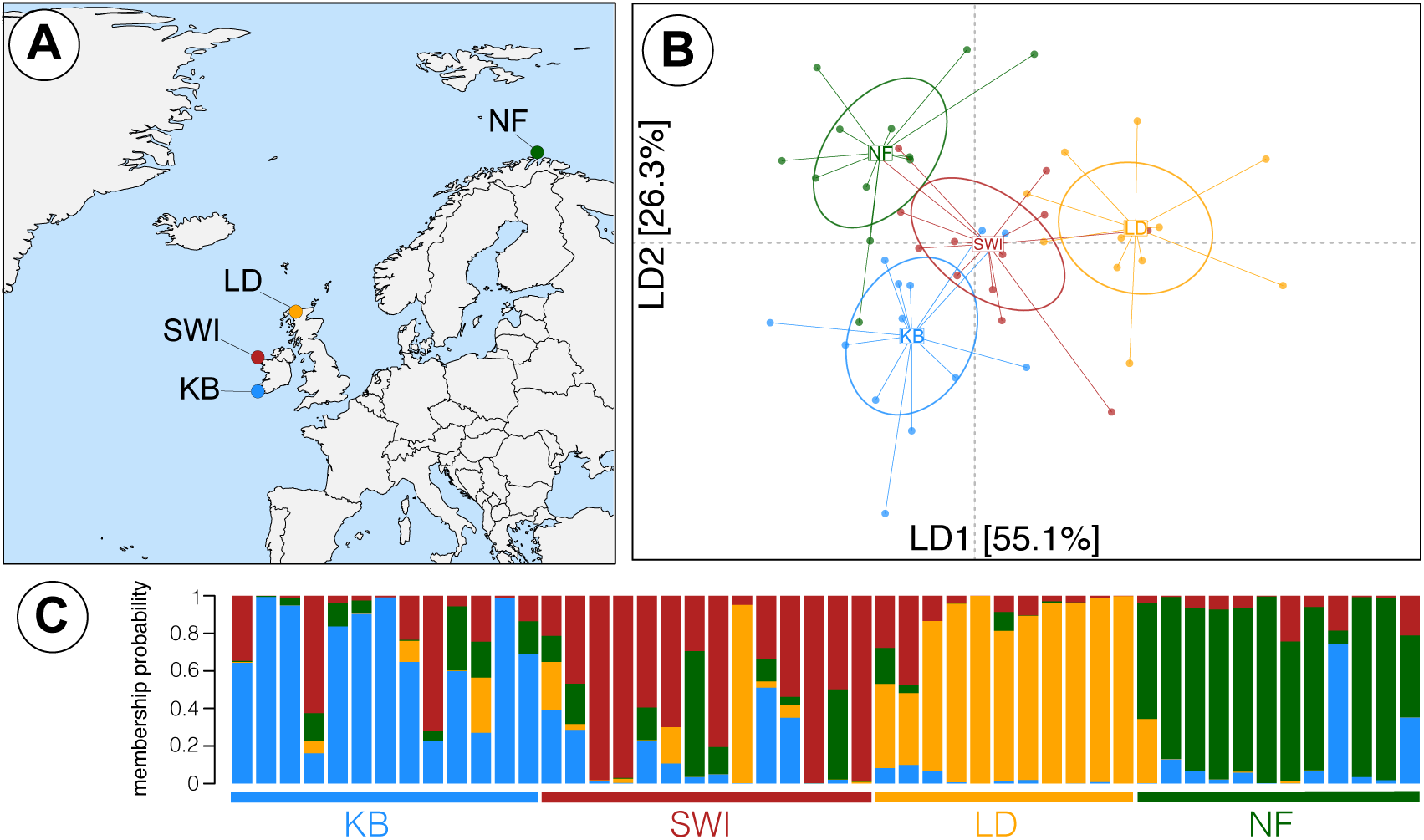
Population structuring in L. salmonis bases on the full SNP dataset. A) Map showing the sampling sites of all four populations across the North-East Atlantic: Finnkirka (NF), Loch Duart (LD), Kenmare Bay (SWI) and Kilkieran Bay (KB). B) DAPC plot of the first and second linear discriminant axis based on the full SNP dataset, explaining a total of 81.4% of the total variation. C) Membership probability plot showing the population assignment probability for each individual.

### IIb-RAD library preparation and sequencing

Library preparation was undertaken as described in Wang et al 2012 (Wang *et al*. 2012). By reference to *in silico* digestion of the reference genome (<u>https://metazoa.ensembl.org/index.html</u>) two enzymes were selected based on potential coverage: AlfI (restrictions site ^5’(10/12)GCA(N)_6_ TGC(12/10)3’^) and CspCI (restriction site ^5’(11/12)CAANNNNNGTGG(12/13)3’^). Digested DNA of each sample was ligated to a pair of partially double-stranded adaptors with compatible and fully degenerated overhangs (5’NNN3’). Finally, the obtained IIb-RAD tags were amplified to introduce a sample-specific 7bp barcode and the Illumina NGS annealing sites using two different pairs of sequencing primers. A 1.8% agarose gel electrophoresis of the PCR products was performed to verify the presence of the expected 150 bp target band (fragment, barcodes and adaptors included). In order to ensure an approximately equimolar contribution of each sample to the library, the concentration of each PCR product was measured from the intensity of the target band in a digital image of the 1.8% agarose gel. We prepared two libraries in total, one for each IIB-REase. The purification of the libraries from high-molecular weight fragments and primer-dimers was achieved first by removing the target band on agarose gel library and eluting them in water overnight; followed by DNA capture with magnetic beads (SPRIselect^®^ Beckman Coulter). The DNA concentration in the purified libraries was quantified with a Qubit^®^ Fluorometer (Invitrogen) and the libraries were combined in one single pool. Two library pools were sequenced, first on a NextSeq 500, then on a MiSeq (Illumina, San Diego, CA, USA) with a single 1x50 bp setup using ‘Version2’ chemistry at Glasgow Polyomics (<u>www.polyomics.gla.ac.uk</u>), which also implemented the read demultiplexing and quality-filtering.

### IIb-RAD data processing

Short reads were aligned to the reference genome in bowtie 2 (Langmead & Salzberg 2012) and SNPs were called using the STACKS v1.42 package with a minimum read depth of 3 (Catchen *et al*. 2013). The *rxstacks* module was used to further screen SNPs and the *population* module filter and export genotypes with a minimum depth of coverage of 6, minimum minor allele frequency of 0.05, maximum observed heterozygosity of 0.5 and present in at least 60% of individuals. To avoid sequence artefacts generated by low complexity in restriction enzyme recognition site, SNPs at positions 12-26 were excluded from the analysis. For those RAD tags that retained diversity after screening for artefacts, only a single SNP per locus was selected for subsequent analysis.

### Population structure analysis and detection of positive selection

Principal components analysis (PCA), discriminant analysis of principal components (DAPC), and population assignment probabilities were calculated in *adegenet* (Jombart & Ahmed 2011). Analyses of molecular variance (AMOVA), Weir and Cockerham estimators of FST, and summary statistics (Ho, He, Gis, π) were calculated in *Genodive* (Meirmans & Van Tienderen 2004). P-values for F_ST_ were FDR adjusted for multiple comparisons using a Benjamini-Hochberg correction in the R-package *p-adjust*. Isolation-by-distance was assessed using a Mantel test implemented in the *vegan* R-package. Loci putatively under positive selection were identified in Lositan (Antao *et al*. 2008) using a FDR < 0.1 significance threshold and localised on the *L. salmonis* linkage map (Glover, K *Pers Comm*) to assess genomic correspondence with a previous population genomic study (Besnier *et al*. 2014). Lositan results were plotted using *ggplot2* in R. Further, we performed a second outlier analysis using BayeScan, as it has a lower type I error rate compared to Lositan (Foll & Gaggiotti 2008)). We ran BayeScan with prior odds of 100 due to the small number of SNPs and detected significant outliers with a FDR threshold of 0.05 and putative outliers with a FDR threshold of 0.1. Finally, we post hoc identified overlapping outlier SNPs between BayeScan and Lositan.

In order to identify genes potentially under positive selection we identified all genes within a 10kb region around each outlier SNP by blasting the sequence against the *L. salmonis* reference genome using the *blastn* function in the EnsemblMetazoa database. We identified all genes within those 10kb regions and when possible determined their function using the *UniprotKB* database.

### Using Random Forest analysis to detect population-discriminatory SNPs

In order to detect SNPs characteristic of each population we employed a machine learning approach using the *randomForest* package in R. Populations were numerically coded and missing data imputed using the *na.roughfix* command. Three independent random forest runs with 100,000 trees each were conducted and checked for convergence between runs by performing pearson correlation between SNP importance values. The resultant dataset (R^2^ > 0.95) was used to select a final dataset for the backwards purging approach. As in Laporte *et al*., all loci with an importance <0 were removed as non-discriminatory (Laporte *et al*. 2016). Backwards purging was performed on the remaining 317 SNPs. As such each random forest run was re-implemented (three independent iterations) and after each run the SNP with the lowest importance was removed until only two SNPs were left. We determined the subset of SNPs with the highest discriminatory power based on the lowest out-of-bag (OOB) error rate and we used this subset for further downstream analysis.

In order to assess the population discriminatory power of the random forest selected SNPs we used the same methods as for the full SNP dataset. First, we performed a PCA and DAPC in *adegenet* in *R* to visualise population structuring and assess the population assignment accuracy. Second, we performed an AMOVA and estimated pairwise Weir and Cockerham’s Fst in *Genodive*. We also identified the overlap between highly discriminatory SNPs and SNPs potentially under positive selection to assess the impact of selection on discriminating *L. salmonis* populations.

## Results

### Bioinformatic processing & summary statistics

Using IIb-RAD sequencing we generated an average of 1,496,567 ± 673,594 reads per individual for 50 individuals from four populations across the North-East Atlantic (Fig. 1a). The final catalogue contained 111,090 RAD tags with an average coverage of 19.6 ± 6.9 per individual, covering 0.34% of the genome. After stringent filtering we retained 1286 SNPs, spanning 787 different reference genome contigs. Genetic diversity, measured as nucleotide diversity (π) and observed heterozygosity (Ho) were similar across populations (Table 1). Tajima’s D did not indicate any signals characteristic of significant population expansion (Fig. S1).

**Table 1:** Summary of sample sizes, mean sequencing coverage per individual and summary statistics, namely observed heterozygosity (Ho), expected heterozygosity (He), inbreeding coefficient (Gis) and genetic diversity (π).

| Population | N | Mean Coverage | $H_o$ | $H_e$ | $G_{is}$ | $\pi$ |
| --- | --- | --- | --- | --- | --- | --- |
| KB | 13 | 19.7 | 0.278 | 0.304 | 0.086 | 0.3025986 |
| SWI | 14 | 18.6 | 0.265 | 0.304 | 0.128 | 0.3019599 |
| LD | 11 | 19.8 | 0.258 | 0.298 | 0.132 | 0.2952346 |
| NF | 12 | 20.2 | 0.267 | 0.312 | 0.143 | 0.3096919 |

### Population structure using the full SNP dataset

In a first approach, we assessed population genetic structure using the full dataset of 1286 SNPs by several different approaches. A PCA did not reveal any population structuring across the entire range, however using pre-defined populations in the DAPC approach revealed a weak population structuring (Fig. 1b & c, Fig. S2). The population assignment probability was on average 0.82 ± 0.10. An AMOVA showed weak but significant population structure (F_sc_ = 0.018, p<0.0001; Table 2). However, based on pairwise F_st_ values only LD and KB were significantly genetically differentiated (F_st_ = 0.01, p<0.0001). No significant isolation-by-distance was detected (R^2^ = -0.35, P = 0.67).

**Table 2:** AMOVA summarising the global population structure in *L. salmonis* populations using the full SNP dataset and only the discriminatory SNPs identified using the random forest appraoch.

|  | Source of Var. | Nested in | % Var | F-stat | F-value | P-value |
| --- | --- | --- | --- | --- | --- | --- |
| Full | Within Ind. | -- | 88.4 | F<sub>it</sub> | 0.116 | -- |
| SNP | Among Ind. | Population | 9.8 | F<sub>is</sub> | 0.1 | p<0.0001 |
| dataset | Among Pop. | -- | 1.8 | F<sub>sc</sub> | 0.018 | p<0.0001 |
|  | Within Ind. | -- | 78.9 | F<sub>it</sub> | 0.211 | -- |
| Discriminatory | Among Ind. | Population | 11.3 | F<sub>is</sub> | 0.125 | p<0.0001 |
| SNPs | Among Pop. | -- | 9.8 | F<sub>sc</sub> | 0.098 | p<0.0001 |

### Using machine learning to define population genetic structure

In order to detect population structuring among populations we utilised a random forest machine learning approach. We detected a subset of 93-101 SNPs that minimised the out-of-bag error rate to 0.1 (compared to 0.76 for the full dataset) and maximised the discriminatory power among populations (Fig. 2A). From this subset, we selected 98 SNPs for further downstream analyses. To assess the power of this subset of 98 highly discriminatory loci to detect significant population structure we performed the same population genetic analysis as was conducted on the full dataset. A PCA performed with the random forest selected subset showed a stronger separation between populations with a weak overlap of 95% confidence-intervals between LD and SWI (Fig. S2). However, the DAPC clearly separated all populations and the population assignment probability recovered was 1, meaning all individuals were correctly assigned to their respective population (Fig. 3). The variance explained among populations increased to 9.8% (from 1.8% with the full dataset) in the AMOVA (F_sc_ = 0.098, p<0.0001). All pairwise comparisons showed highly significant Fst-values (range = 0.081 – 0.096), confirming the significant discriminatory power of the random forest detected SNP subset.

**Figure 2:**
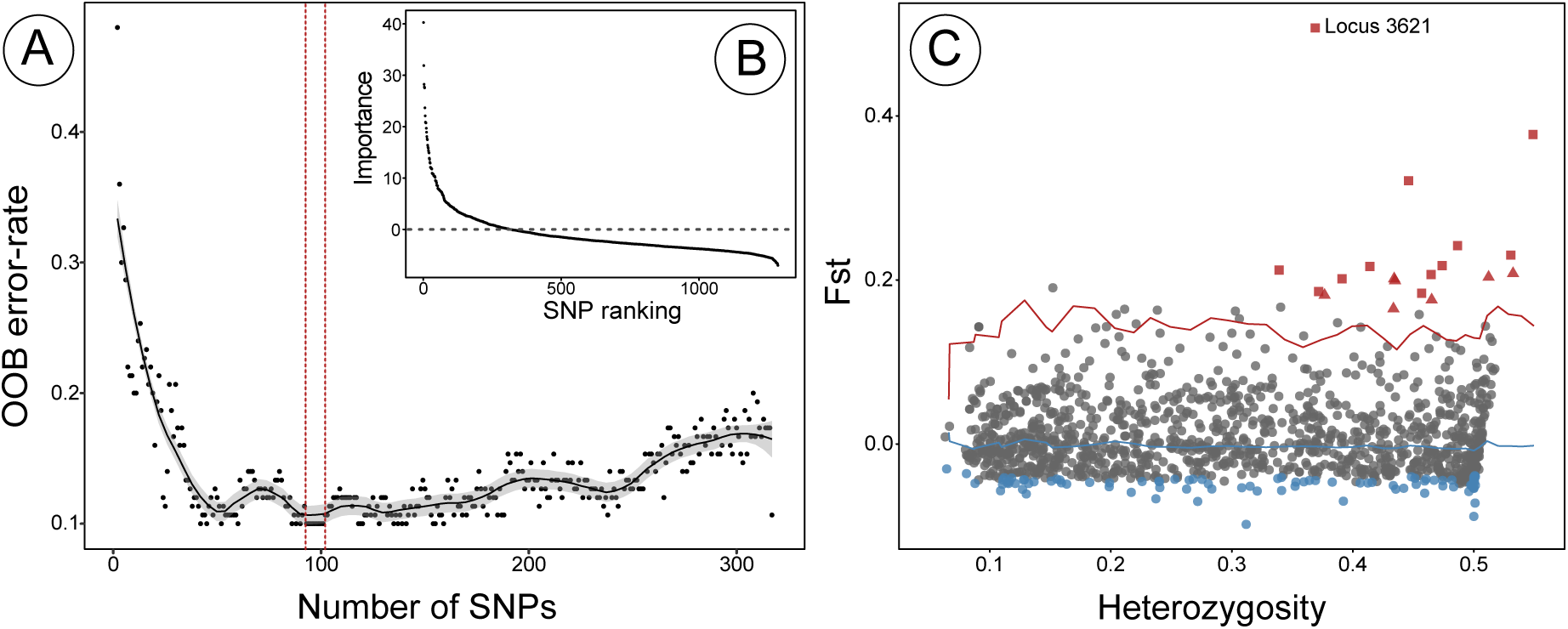
Detecting discriminatory loci using random forest and signals of selection. A) Plot showing the results of the backwards purging approach, with the number of SNPs per subset plotted against the out-of-bag (OOB) error rate for each subset. The black line shows the smoothed estimates with 95% confidence-intervals (grey area). The two red dotted lines show the range of subsets (93-101 SNPs) with the lowest OOB error rate. B) The inset shows the initial distribution of scaled importance values for each SNP before the backwards purging. The grey dotted line shows the importance threshold for the subset of SNPs used for backwards purging. C) F_ST_ outlier analysis results showing individual SNP loci and 5% (blue line) and 95% (red line) confidence intervals. Outlier loci potentially under positive selection are in plotted in red and those potentially under balancing selection in blue. Squares mark F_ST_ outlier loci that were also detected as highly discriminatory using random forest and triangles those that are not shared. The significant outlier detected using BayeScan is labelled with ‘Locus 3621’.

**Figure 3:**
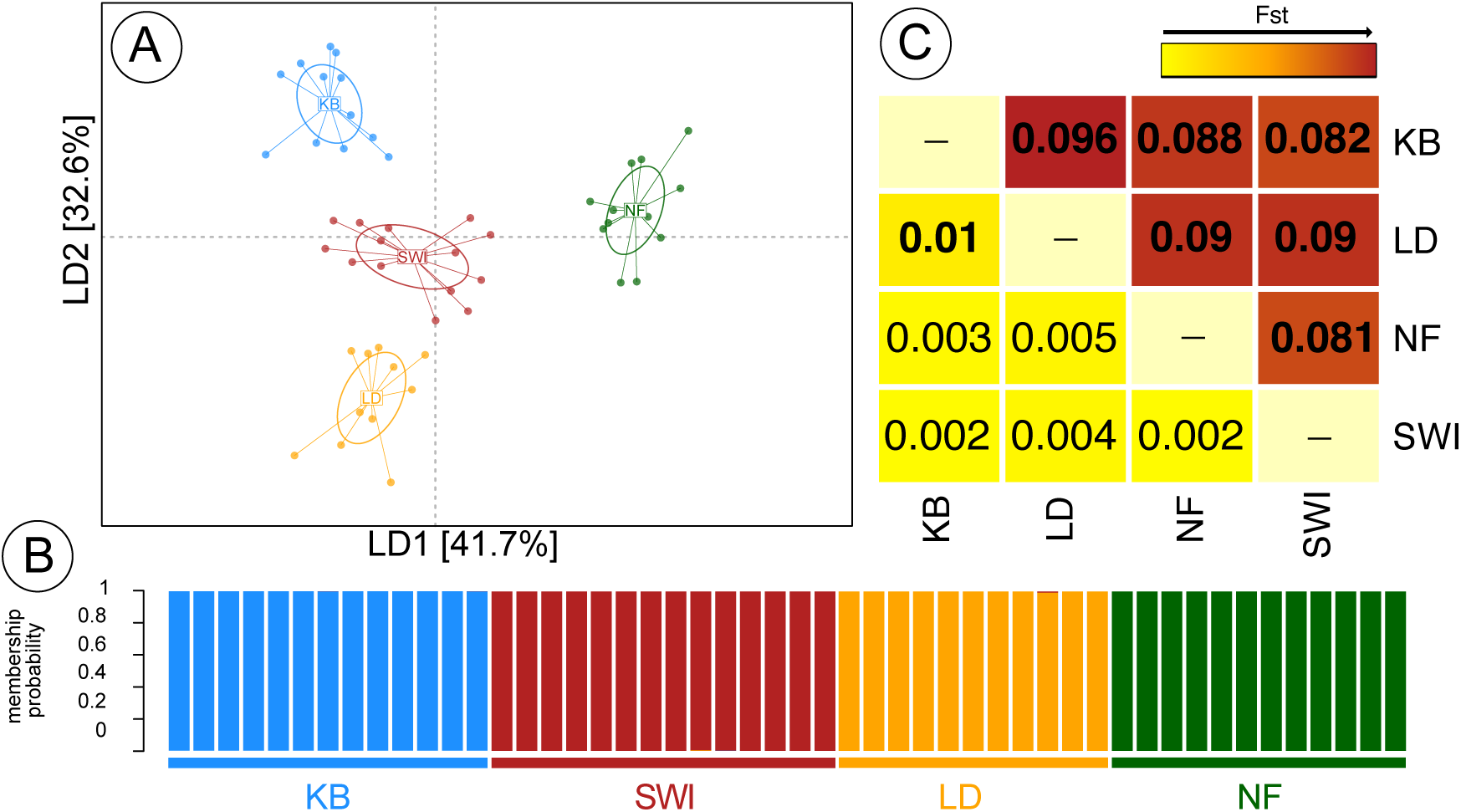
Population structure and population assignment in *L. salmonis* using discriminatory random forest loci. A) DAPC plot of the first and second linear discriminant axis based on 98 highly discriminatory SNPs, explaining a total of 74.3% of the total variation. B) Membership probability plot showing the population assignment probability for each individual. Each individual was correctly assigned to its sampling site. C) Heatmap showing pairwise Fst between sampling sites based on the full SNP dataset (below diagonal) and based on the highly discriminatory SNP subset (above diagonal). Significant Fst values (inside each square) with P < 0.05 are highlighted in bold.

### Population discriminating SNPs and selection

One factor that might explain the strong discrimination of sea louse populations using the subset of random forest-selected SNPs would be divergent selection pressures, such as adaptation to different drug treatments, or local adaptation to natural environmental factors. Therefore, we performed two different tests for selecting SNPs under significant positive selection. An FDist approach implemented in Lositan detected 19 SNPs under strong positive selection (FDR < 0.1) with an average F_st_ of 0.233 ± 0.083 between populations (Fig. 2C, Table 3). Eleven out of all 19 SNPs under positive selection are located on previously defined linkage groups 1 and 14, seven and four respectively (Besnier *et al*. 2014). The remaining SNPs are either located on linkage groups 4, 6 and 7 (two, two and one respectively) or could not be assigned to a linkage group. We further detected 46 SNPs under balancing selection (FDR < 0.1; p < 0.02).

**Table 3:** Outlier SNPs identified using the different approaches (Lositan, BayeScan and Random Forest) and annotation. RF stands for random forest, meaning SNPs that have been detected using the random forest approach. ‘Unchar.’ stands for uncharacterized genes.

| Locus ID | Contig_position | LG | Fst (Lositan) | Lositan | BayeScan | RF | Annotation |
| --- | --- | --- | --- | --- | --- | --- | --- |
| 38173 | lsalatl2s740_42780 | 4 | 0.320968 | Yes | Putative | Yes | -- |
| 3621 | lsalatl2s1185_140991 | 1 | 0.50726 | Yes | Yes | Yes | -- |
| 40396 | lsalatl2s80_965936 | 1 | 0.20663 | Yes | No | Yes | PSA2 |
| 41679 | lsalatl2s85_1109389 | 4 | 0.181416 | Yes | No | No | -- |
| 42860 | lsalatl2s907_144760 | -- | 0.203467 | Yes | No | No | -- |
| 4355 | lsalatl2s122_618061 | 7 | 0.207882 | Yes | No | No | -- |
| 6832 | lsalatl2s139_1380660 | 1 | 0.199199 | Yes | No | No | -- |
| 8287 | lsalatl2s14_555303 | 1 | 0.377272 | Yes | Putative | Yes | unchar. |
| 8241 | lsalatl2s14_1020918 | 1 | 0.217428 | Yes | No | Yes | unchar. |
| 9674 | lsalatl2s163_163880 | 14 | 0.201546 | Yes | No | No | -- |
| 15099 | lsalatl2s228_333839 | 1 | 0.241913 | Yes | No | Yes | -- |
| 1623 | lsalatl2s10843_736 | -- | 0.212014 | Yes | No | Yes | -- |
| 21928 | lsalatl2s3387_1782 | -- | 0.185878 | Yes | No | Yes | -- |
| 25024 | lsalatl2s39_920686 | 6 | 0.216516 | Yes | No | Yes | -- |
| 26383 | lsalatl2s429_103294 | 14 | 0.175808 | Yes | No | No | unchar. |
| 29942 | lsalatl2s514_325267 | 14 | 0.164932 | Yes | No | No | -- |
| 30716 | lsalatl2s535_184954 | 14 | 0.230334 | Yes | No | Yes | -- |
| 2652 | lsalatl2s1135_117353 | 1 | 0.183753 | Yes | No | Yes | -- |
| 30805 | lsalatl2s538_341294 | 6 | 0.20138 | Yes | No | Yes | unchar. |

An analysis in BayeScan detected only one SNP (FDR < 0.01) under significant positive selection and two more putatively under selection (FDR < 0.05). All three of these SNPs were also detected to be under selection by Lositan and the significant one was also the top outlier in Lositan and located on linkage group 1 (F_st_ = 0.507). The other two putative SNPs in BayeScan were also highly significant in Lositan (p < 0.001) and were located on linkage group 1 and 4.

To detect how selection influences the genetic discrimination of populations we identified the amount of overlap between the 98 SNPs detected by random forest and all Lositan SNPs under significant positive selection. 63.2% of loci (12 out of 19 loci) detected to be under positive selection using Lositan were also identified being highly discriminatory between populations using random forest. Locus 3621, which was also identified using BayeScan, had the highest importance in the random forest analysis, suggesting that strong local adaptation and selection distinguishes sea louse populations.

### Annotation of outlier SNPs

In order to identify specific genes potentially involved in local adaptation and are under positive selection in sea lice, we explored these regions in the annotated *L. salmonis* genome. Five of the 19 SNPs were in regions contained annotated genes within 10kb, but only one of the annotated genes has been characterized. Two of the contigs with annotated genes were on linkage group 01, two on linkage group 14 and one on linkage group 6. Contig LSalAtl2s80 (linkage group 01) contained the characterized gene *PSA2*, which codes for the proteasome subunit alpha type protein.

### Discussion

In this study, we used a IIb-RAD sequencing approach coupled with advanced and sensitive population genetic analyses to genetically ‘fingerprint’ *L. salmonis* populations in the North-East Atlantic and to detect signatures of selection. We were able to achieve this using a relatively small (n=50) number of individuals genotyped across only a limited portion of the genome (c.2.3Mbp = 0.34%). An important set of discriminatory loci was identified against a background of high genetic connectivity via a random forest machine-learning algorithm and these can be exploited to distinguish between nearby sea louse populations. A high degree of overlap between loci under positive selection using genome-scan approaches and loci with high discriminatory power from random forest analysis was also observed.

Sea lice are known to disperse rapidly among aquaculture sites as part of the larval zooplankton as well as via the movements of migratory (*Salmo salar)* or resident (*Salmo trutta*) anadromous salmonids (Boxaspen 2006). Previous population genetic studies were consistent with such high levels of dispersal (Glover *et al*. 2011; Nolan & Powell 2009; Todd *et al*. 2004), finding no significant genetic differentiation in our study region when utilizing a set of neutral microsatellite loci. Inclusion of putatively non-neutral loci can improve population discrimination across the Atlantic (e.g. (Glover *et al*. 2011)). However, the same studies could not distinguish populations on a small geographic scale as our data and approach suggest is possible.

More recent genome wide analysis of SNP variation in *L. salmonis* has to date been consistent with the lack of genetic structure that was found using classic markers such as microsatellite loci (Besnier *et al*. 2014). As with our dataset, correlation with geographic distances is not a feature of the genetic variation observed even with such genome-wide information. We found global F_ST_-values based on all loci to be significant but low (0.018), in agreement with patterns that have been found previously (Besnier *et al*. 2014). The use of anti-parastic drugs has been shown to be a strong selective pressure in sea lice and several genomic regions under selection have been linked to drug treatment (Besnier *et al*. 2014). In particular linkage groups 1 and 5 in the study showed evidence of selective sweeps, with the same region on linkage group 5 being implicated in drug resistance in a QTL analysis (Besnier *et al*. 2014). Other linkage groups, such as 14 also showed signal of positive selection in that study (Besnier *et al*. 2014). Our study similarly found that 11 out of the 19 outlier loci we identified also lay in linkage groups 1 & 14, which represents an important independent validation. Spatio-temporal variation in treatment regimes, such as rotations of different drugs or the alternative use of warm-water or freshwater treatments (Ljungfeldt *et al*. 2016), may drive the heterogeneity observed in our and previous studies. This is partly as a result of cost, perceived efficacy, as well as different regulatory conditions in the countries concerned. Even though spatio-temporal variation in drug resistance is likely to be the strongest driver of differential selection among populations, local environmental conditions can constitute further selective pressures driving allele frequency differences among populations. Local environmental variables such as temperature (e.g. (Samsing *et al*. 2016)) and salinity (e.g. (Bricknell *et al*. 2006)), for example, can have profound effects on sea louse survival and development. Furthermore, a combination of drug treatment and increased host density is shown to influence the evolution of reproductive and life history traits (Mennerat *et al*. 2017), as well as virulence in sea lice (Mennerat *et al*. 2012) among different populations. However, such local adaptation is most likely linked to subtle allele frequency differences, compared to strong selective sweeps caused by drug treatments, as the selective pressure is comparably low. The combination of a few outlier loci under strong positive selection and a wide range of loci showing subtle allele frequency differences fits the expected pattern. Independent of the cause for allele frequency differences among populations, we show that a random forest machine learning approach can be used to cost-effectively distinguish even near-by sea louse populations, even with a low number of samples and genotyping density.

The use of (historical) samples from the same site at different time points, differing in treatment regimes, could be used to disentangle the effects of drug regime and local adaption on allele frequency differences among populations and signatures of selection. Genome-scale population genetic profiling, alongside robust phenotyping, may also eventually reveal the genetic architecture underlying drug resistance and local adaptation. Here we have identified signals of selection across the genome, including markers closely associated with functional genes (e.g. *PSA2)*. The association of genomic response to selection, natural environmental conditions, and drug treatment profiles will be important considerations for future work.

Tools to enable parasite traceability and molecular epidemiology are an important requirement for rational sea louse control. Hydrographic modelling has been successfully deployed to understand short-range dispersal *L. salmonis* between farms and have been used to evaluate optimal treatment strategies (Gettinby *et al*. 2011; Salama *et al*. 2013). Such model predictions can be biologically ‘truthed’ using planktonic trawls and strategically placed ‘sentinel’ fish that can infer the geographic scales of dispersal, as it has been done in one of the study areas, Kilkieran Bay (KB), (Jackson *et al*. 2012). However, biological (or genetic) confirmation of larger scale dispersal models (i.e. between lochs (=fjord) and loch systems) within and across regions is also required to assess long-range re-infestation risks for aquaculture sites. Such a strategy is of particular relevance in the light of an increasing control focus on loch-wide fallowing practices (Torrissen *et al*. 2013). Furthermore, integration of genetic connectivity data with hydrographic larval dispersal models – so called ‘seascape genetics’ (e.g.(Riginos & Liggins 2013)) - is likely to be more fruitful in defining any spatial-genetic correlations than crude map distances and represents an interesting further avenue for study. In this context, our data show that it may be possible to genetically ‘fingerprint’ louse populations in nearby regions to understand connectivity between them and provide a valuable tool for disease surveillance.

## Acknowledgements

Many thanks to Gary Carvahlo and Simon Creer, University of Wales, Bangor for providing logistical support in the early stages of this work. Thanks also to the Glasgow Polyomics team, Julie Galbraith, Graham Hamilton and Pawel Herzyk for assisting with library preparation and sequencing. Finally, thanks to Kevin Glover and Rasmus Skern at the Sea Louse Research Centre for generously providing access to the linkage maps and drafts of the *L. salmonis* genome.

## Data Archiving Statement

All reads will be submitted to the NCBI Short Read Archive (Accession XXXX) and will be made available upon acceptance.

## Supplementary material

**Figure S1**: Violin plots of Tajima’s D for each sea louse population.

**Figure S2**: Principal component analysis for the full SNP dataset and random forest candidate SNPs.

